# Different obesity phenotypes correlated with the PI3K-AKT signaling pathway in tree shrews

**DOI:** 10.1101/361063

**Authors:** Yuanyuan Han, Huatang Zhang, Jiejie Dai, Yang Chen

## Abstract

**Objective:** To identify the meaningful co-expressed gene network in mRNA extracted from adipose tissue representing different obesity phenotypes in a new tree shrew model. Furthermore, to locate the potential drug target based on analyzing the possible pathways and hub genes responsible for obesity.

**Methods:** Ten tree shrews were selected from F1 populations and divided into 3 groups based on their Lee’s index for mRNA sequencing. We identified clusters of highly correlated genes (modules) by weighted gene co-expression network analysis (WGCNA).

**Results:** Three modules were strongly correlated with not less than one obesity phenotype (associations ranging from −0.94 to 0.85, P < 0.01). The ribosome, lysosome and ubiquitin-mediated proteolysis pathways were the most enriched pathways of the blue (including 481 genes), brown (389 genes) and turquoise (1781 genes) modules, respectively. The hub genes were determined, including UBA52 in the blue module, AKT1 in the brown module and LRRK2 in the turquoise module.

**Conclusions:** We characterized different obesity phenotypes in tree shrews for the first time. The most represented gene AKT1 indicated the vital role of the PI3K-AKT pathway through interactions with downstream pathways and differentially expressed genes (DEGs). The work will help provide new insight into the prevention and treatment strategies of obesity.

## Background

The tree shrew (*Tupaia belangeri*) is a prospective laboratory animal that has a closer genetic association with primates than with rodents^1,2^. Additionally, other advantages including easy maintenance, small body size, and rapid reproduction make the tree shrew an excellent model animal. Many studies use the tree shrew for fundamental biological mechanisms ^3–5^, modeling human diseases ^6–11^, organism responses ^12,13^ and molecular evolution^14,15^. The tree shrew has been proposed as a substitute choice to primates in biomedical research. The available annotated genome data provide a solid foundation to analyze the genes of tree shrew at the transcriptomic level^16,17^.

Obesity is not only a human problem^18^. Researchers first noticed the obese phenotype in artificial breeding tree shrews in 2012^19^. According to X. Wu et al., the Lee’s index score threshold of 290 distinguishes whether a tree shrew is obese. Thus, tree shrews are considered obese with a score above 290 and non-obese with a score below 290 (whereas in rats, obesity starts at a Lee’s index of 310, no universal definition of obesity is established for tree shrews). The situation is more difficult because the average weight of tree shrews has increased in the latest years. Similar to humans, increasingly caloric diets and limitation of exercise in breeding cages are the primary causes of tree shrew obesity. However, increasingly fat tree shrews in controlled lab settings are surprising. Thus, what is the mechanism underlying this tree shrew obesity? What is the difference between obese and non-obese shrews? Are the same processes and key factors shared by human and tree shrew obesity? Identification of gene networks in mRNA extracted from relevant tissue samples from different obesity phenotypes of tree shrews may provide an entry point to answer these questions.

Obesity is an over-nutrition condition, which harms systemic metabolic balance and evokes stress^20^. Chronic inflammation in white adipose tissue (WAT) is a characteristic of obesity pathophysiology and is highly associated with other severe diseases^21^. The initiation and exacerbation of chronic inflammation primarily occurs in WAT^22–24^. Therefore, considering the central role of WAT in energy homeostasis^25^, we used subcutaneous adipose tissue from the tree shrew model of spontaneous obesity to study the mRNAs of three extreme groups. We performed the WGCNA^26^ method using RNA-sequencing (RNA-Seq) data. We aimed to elucidate transcriptomic mechanisms of obesity in tree shrews and to shed light on the mechanism of human obesity by detecting mRNAs and pathways involved in the pathogenesis of obesity^27^.

In this study, we report spontaneous physiological changes and obesity-related health conditions, including increased abdominal fat, BMI and alterations in various serum chemistry parameters, in the tree shrew similar to those observed in humans. We further categorize the animals into three groups, lean and moderate and severe obesity, based on body weight distribution and Lee’s index among the entire population, as is performed in humans. Elucidation of the transcriptomic factors that contribute to excessive abdominal fatness in tree shrews in a relatively short time could advance our understanding of the mechanism underlying obesity and similar human metabolic disease conditions.

## Methods

### Profile of tree shrew obesity

Three hundred and twenty-nine F1 tree shrews (168 males and 161 females) were bred in the Centre of Tree Shrew Germplasm Resources, Institute of Medical Biology, Chinese Academy of Medical Science and Peking Union Medical College (IMBCAMS). The age of the 329 tree shrews ranged from 1.5 to 4.5 years. All methods were conducted in accordance with ethical and relevant guidelines and regulations. The institutional Animal Care & Welfare Committee of IMBCAMS^28^ approved all experimental protocols.

The body weight and Lee’s index [(body weight (g)* 1000)^(1/3)/body length (cm)*100] were determined as before^19^. Based on the normal distribution curve of Lee’s index, tree shrews were categorized in three groups: lean (Lee’s index ≈mean + SD), moderate fat (Lee’s index >mean + 2 SD) and severe fat (Lee’s index >mean + 4 SD).

We chose 10 animals from these 3 groups and used the subcutaneous adipose tissue for RNA-Seq: three severe fat (O), three moderate fat (M) and four lean (L) tree shrews.

### Phenotypic characterization of the selected tree shrews

Data acquisition on all 329 tree shrews was completed within a 1-week time in April 2017. All tree shrews were fasted for 12 h before measuring. Each tree shrew was caught in a pocket, and the body weight/length were measured twice. Biochemical parameters and related data on the 10 selected tree shrews were determined as described previously^19^. An Aquillon ONE 320-row helical CT scanner (Toshiba, Tokyo, Japan) was used to determine fat volume in vivo. The results were considered significant at P < 0.05.

### Sample collection and preparation

The mRNA was extracted using TRIzol^™^ Reagent (Thermofisher, MA, USA). Extracted RNA purity was assessed by a NanoPhotometer^®^ spectrophotometer (IMPLEN, CA, USA). RNA concentration was measured using a Qubit^®^ RNA Assay Kit in a Qubit^®^ 2.0 Fluorometer (Life Technologies, CA, USA). RNA integrity was assessed using an RNA Nano 6000 Assay Kit of a Bioanalyzer 2100 system (Agilent Technologies, CA, USA). Sequencing libraries were generated using a NEBNext^®^ UltraTM RNA Library Prep Kit for Illumina^®^ (NEB, USA) following the manufacturer’s recommendations. The clustering of the index-coded samples was performed on a cBot Cluster Generation System using a TruSeq PE Cluster Kit v3-cBot-HS (Illumina) according to the manufacturer’s instructions. After cluster generation, the library preparations were sequenced on an Illumina Hiseq platform and 125 bp/150 bp paired-end reads were generated. Before alignment, clean data (clean reads) were obtained by removing reads containing adapters, reads containing ploy-N and low quality reads from raw data. The clean reads were aligned to the reference genome TupChi_1.0 using STAR (v2.5.1b) with the method of Maximal Mappable Prefix (MMP).

### Quantification of gene expression level and (DEG) expression analysis

The number of reads aligned to every gene was estimated by HTSeq v0.6.0. Subsequently, the FPKM of all genes was determined, resulting in an average of 10,838 genes per sample. The DEG analysis between each two groups was conducted using the DESeq2 R package (1.10.1). DESeq2 provides statistical routines for determining DEGs from sequencing data that rely on the negative binomial distribution. The resulting P-values were adjusted using Benjamini and Hochberg’s approach to control the false discovery rate. Genes with an adjusted P-value using Benjamini and Hochberg’s method < 0.05 were designated DEGs.

### WGCNA

We reduced the data set further before network construction, and genes were selected based on variation (SD > 0.25), leading to 7,115 genes. We conducted WGCNA as mentioned previously^27^. Gene ontology (GO) and the Kyoto Encyclopedia of Genes and Genomes (KEGG) analysis.

GO and KEGG were used to interpret gene data in each module^28,29^. The DAVID database (https://david.ncifcrf.gov/) is a combined data source for annotation, visualization and integrated GO/KEGG analysis for a set of genes. Functional and pathways analyses were conducted using this database to identify the gene function and most enriched pathways of DEGs. The cut-off criterion was an adjusted P value < 0.05. We selectively showed the top 10 significantly enriched GO terms and showed all the significantly enriched pathways.

Construction and analysis of the protein-protein interaction network

DEGs in each module were mapped to the PPI data via the Search Tool for the Retrieval of Interacting Genes (STRING) database v.9.1 (http://www.stringdb.org/) to demonstrate potential protein-PPI networks^31,32^. The STRING database applies a meta-method to assess PPI relationships and identifies functional or physical links between proteins. The extended network was built based on the minimum required interaction score of 0.9, which indicates that only interactions with the highest confidence score were selected. Cytoscape is a useful software tool for visually exploiting biomolecular interaction networks. The DEGs in each module were mapped to STRING at first and then visualized using Cytoscape. The criterion for screening hub genes was based on the node degree. Phylogenetic analysis was conducted using previously described methods^30^. The three-dimensional structure of the protein was determined by SWISS-MODEL and presented by Pymol as mentioned previously^28^.

### Hub gene analysis and validation

Hub genes are highly interconnected with one another in a module and have evolved in their function. We identified the hub genes based on both their module membership (>0.9) and their node degree in the PPI network. We conducted linear regression analyses between relative expression of AKT1 and obesity phenotypes (including body weight, Lee’s index, blood sugar and triglycerides) to validate the selected hub gene in tree shrew obesity.

## Results

### Weight and Lee’s index of surveyed tree shrews

The body size of the animals in the severe obese group was significantly different from that of their lean counterparts (Figure 1A). Body length and body weight measurements (Figure 1B and 1C) of the 329 breeding tree shrews were conducted within a 1-week period in April 2017. Tree shrew age was between 1.5 and 4.5 years (Figure 1D). The average body weight of the 329 tree shrews was 148.47 ± 15.09 g, and the average Lee’s index was 280.67 ± 15.09. Although the body weight and Lee’s index were variable, the data followed a Gaussian distribution (Figure 1E). However, Pearson’s correlation analysis suggested that body weight (r = 0.27, P <0.001, Figure 1F) and Lee’s index (r = 0.41, P <0.001, Figure 1G) were positively correlated with age, which was equal to feeding time. The body weight and Lee’s index were significantly higher than those previously reported for wild tree shrews (128.66 ± 0.66 g and 274.46 ± 8.16, respectively).

**Figure 1.**
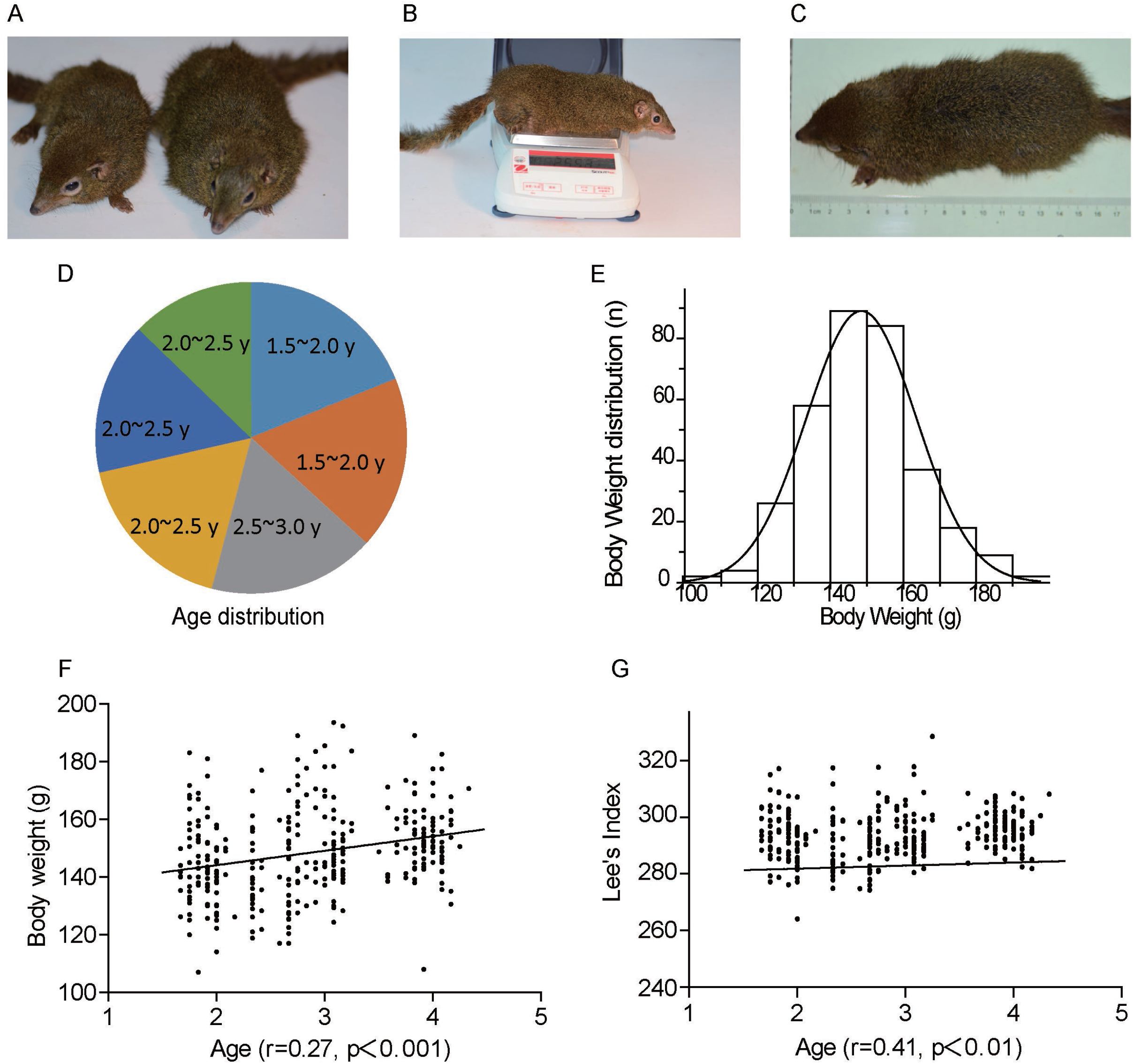
Survey of body weight and Lee’s index of tree shrews. A) Obese tree shrew with lean counterpart. B) Body length measurement. C) Body weight measurement. D) Age distribution. E) Body weight distribution. F) Correlation between body weight concentration and age in tree shrews. H) Correlation between Lee’s index and age in tree shrews.

### Animals selected on the basis of body weight, body length and Lee’ index

The normally distributed body weight and Lee’s index for the entire population clearly showed that the degree of obesity varied widely in these animals. Based on the population distribution of body weight and Lee’s index, we categorized the animals into three groups: severe obese, lean and moderate obese. On the basis of the above criteria, we selected 10 tree shrews. The body weight, Lee’s index and abdominal circumference were significantly different among the three groups. The blood glucose concentration after fasting in severe and moderate obesity groups was significantly higher than that in the lean group. The level of serum HbA1c in the moderate obese tree shrews (7.83 ± 4.56) was the highest of the three groups. In spite of the numeral differences in fasting blood glucose and serum HbA1c concentrations, no statistically significant differences in these parameters were detected between any two groups. CT scan analysis showed 3 and 6-fold increases in total fat volume in the moderate and severe obese tree shrews, respectively, compared with the volume in the lean group. These results are typical of obese humans and other commonly utilized models of obesity.

Adipose tissues from four lean (L), three moderate obese (M) and three severe obese animals (O) were obtained for RNA-Seq for a total of 10 samples. Descriptive statistics for each obesity phenotype group are shown in Sup table 1. The average age of the selected tree shrews was between 2 and 4 years at slaughter.

### Differential gene/transcript expression

A total of 1679 DEGs were identified between M and L, 6961 between M and O and 6376 between O and L, and we selected these DEGs (fold change >2) for subsequent bioinformatics analysis. The heat map of the Pearson’s correlation matrix for the DEGs among the three extreme groups indicated that the three groups also differed at the transcriptomic level, consistent with the obesity phenotypes (Figure 2A). DEGs among the three groups are shown together in Figure 2B. The obtained clean data was submitted to NCBI Sequence Read Archive database (submission ID: PRJNA310673).

**Figure 2.**
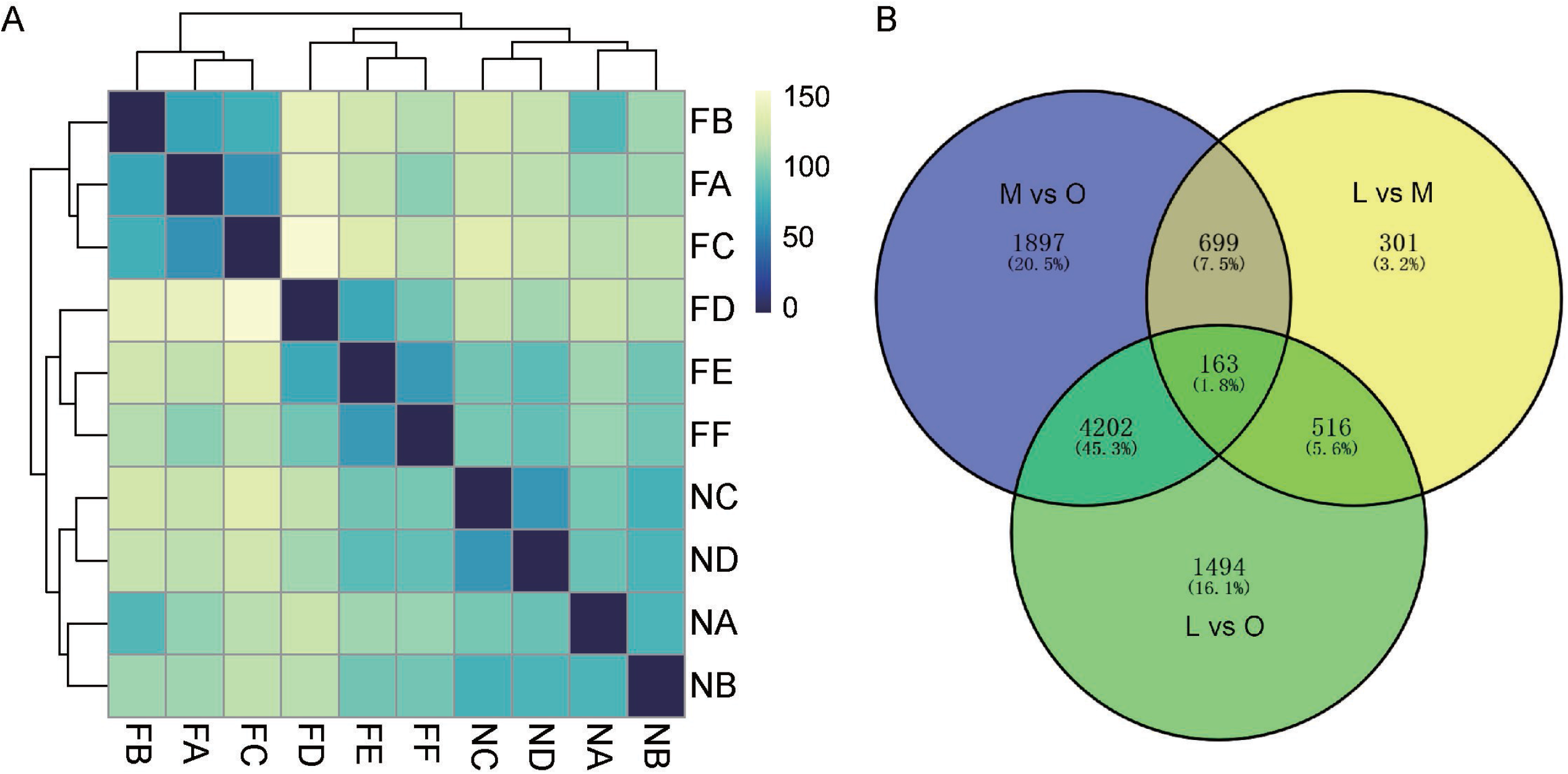
Differentially expressed genes (DEGs). A) Pearson correlation matrix of transcriptome data. B) Venn diagram of DEGs among 3 groups.

### WGCNA

We employed WGCNA to evaluate the RNA-Seq data. The WGCNA assumes that highly co-expressed genes work cooperatively, contributing to the corresponding phenotype. The network was constructed using 7,115 count genes. Strongly co-expressed genes in clusters (modules) were found and assigned to module colors (Figure 3A). In all, we found 3 modules with each covering at least 300 genes.

**Figure 3.**
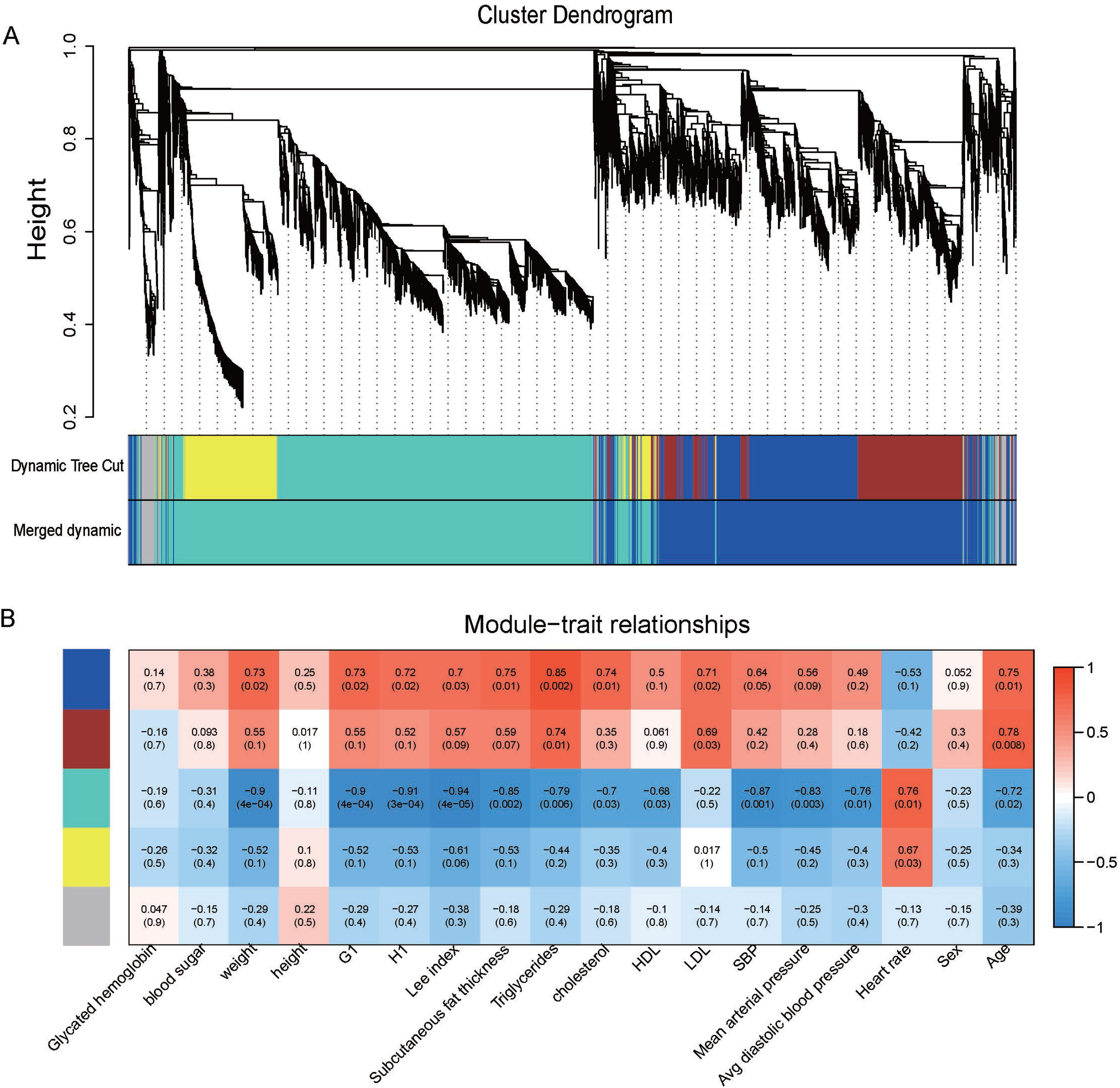
WGCNA results. A) Average linkage clustering tree (dendrogram) defined by WGCNA representing the co-expression modules. Branches of the dendrogram correspond to modules labeled with different colors below the dendrogram. B) MTRs and matching P-values between the modules and their related traits. The discovered modules are on the y-axis, and the discovered traits are on the x-axis. Other color rows showed the MTRs’ degree of correlation. Red indicates a strong positive association; blue indicates a strong negative correlation.

An eigengene was calculated, representing each module to explain the gene expression variation. The module-trait relationships (MTRs) were measured by linking the eigengene to the traits. Then, MTRs were used to select relevant modules for further analysis (Figure 3B). Based on the criteria of both MTRs>0.5 and their gene intra-modular connectivity mentioned previously, we selected 3 modules for further annotation: the blue module (481 genes), the brown module (389 genes) and the turquoise module (1781 genes).

### Functional enrichment of DEGs in modules

We found enriched GO terms and KEGG pathways in the 3 modules. P-values were adjusted using the Benjamini-Hochberg (BH) correction. We selectively showed the top 10 significant GO biological process terms and all the significant KEGG pathways (Sup table 2).

The most interesting and relevant module was the blue module, which showed high MTRs with obesity-related traits, such as triglycerides (0.85), subcutaneous fat thickness (0.75) and age (0.75). The genes in the blue module were highly expressed in obese tree shrews and showed low expression in lean tree shrews. This module showed significant translation and metabolism-related GO terms and KEGG pathways after BH correction. The most enriched GO term was a multi-organism metabolic process (Padj =2.93E-39). The highest enriched pathways were related to pyrimidine, purine, and cysteine metabolism.

Enrichment analysis of KEGG pathways found five pathways at the P < 0.05 level. The most notable were the ‘ribosome’, ‘spliceosome’ and ‘RNA polymerase’ pathways. The enriched pathways and contained genes are collectively presented in Sup table 3. The fourth significant KEGG pathway was ‘protein processing in the endoplasmic reticulum (ER)’, with protein translation, folding, sorting and degradation occurring in the ER. The ER can also coordinate various cellular processes through unfolded protein response (UPR) signaling. Obesity murine models are accompanied by chronic UPR activation in liver and/or adipose tissues^33^.

The brown module (eigengene), exhibited an overrepresentation of immune stress relevant GO terms and KEGG pathways, e.g., 83 genes enriched in immune system processes (Padj =6.75E-12) and 102 genes enriched in the regulation of response to stimulus (Padj =7.36E-07) in GO biological processes. These expression profiles are consistent with previous studies. Much previous work demonstrates chronic inflammation in hypertrophied adipocytes and increased levels of proinflammatory cytokines ^20^. Macrophages play a key role in the initiation and exacerbation of the inflammatory state in obesity^34^. Phagocytosis is a primary mechanism used to remove pathogens and cell debris by fusing phagosome and lysosomes to form a phagolysosome in macrophages. As expected for the 389 genes whose module membership was greater than 0.9, we observed the phagocytosis-related GO pathways, e.g., ‘endocytosis’ (Padj =0.001) and ‘phagocytosis’ (Padj =0.001). The most significant pathway were also phagocytosis-related. A significant immune-related KEGG pathway was ‘natural killer cell-mediated cytotoxicity’ (Padj =0.0077). High BMI in tree shrews is associated with ‘increased activation of peripheral NK cells’. A recent study reported that the activated NK cells in obesity patients were hard to degranulate or to produce signal cytokines. Thus, constant stimulation of NK cells may lead to an abnormal response, which could make obese individuals more susceptible to infectious diseases^35^.

The cellular responses to stimulus and inflammation can inhibit or activate one another. The immune system regulates the cellular stress through signaling proteins and pathways, e.g., ‘regulation of response to stimulus’ (GO:0048583, Padj =1.76E-11), ‘regulation of immune response-regulating cell surface receptor signaling pathway’ (GO:0002768, Padj =6.78E-11), ultimately causing metabolic changes and subsequent altered insulin sensitivity.

The turquoise module showed a strong negative correlation with Lee’s index (−0.94). The module also showed significant signaling-related GO terms and KEGG pathways. We identified 13 pathways at the level P < 0.05. Notable among the pathways were ‘ubiquitin-mediated proteolysis’ (Padj =1.60E-05), ‘phosphatidylinositol signaling system’ (Padj =0.00562), ‘TGF-beta signaling pathway’ (Padj =0.012), and ‘mTOR signaling pathway’ (Padj =0.017)’.

To further elucidate the enriched trend of each module, we selected the top 50 eigengenes in each module to demonstrate the expression levels of the genes therein. The genes in the blue module were highly expressed in severe obesity, moderately expressed in moderate obesity and expressed at low levels in lean animals (Figure 4A). The genes in the brown module were highly expressed in both severe obesity and moderate obesity and expressed in low levels in lean animals (Figure 5A). The GO/KEGG pathways of the top 50 genes in the blue (Figure 4B) and brown modules (Figure 5B) were further analyzed to show the most relevant GO terms and pathways.

**Figure 4.**
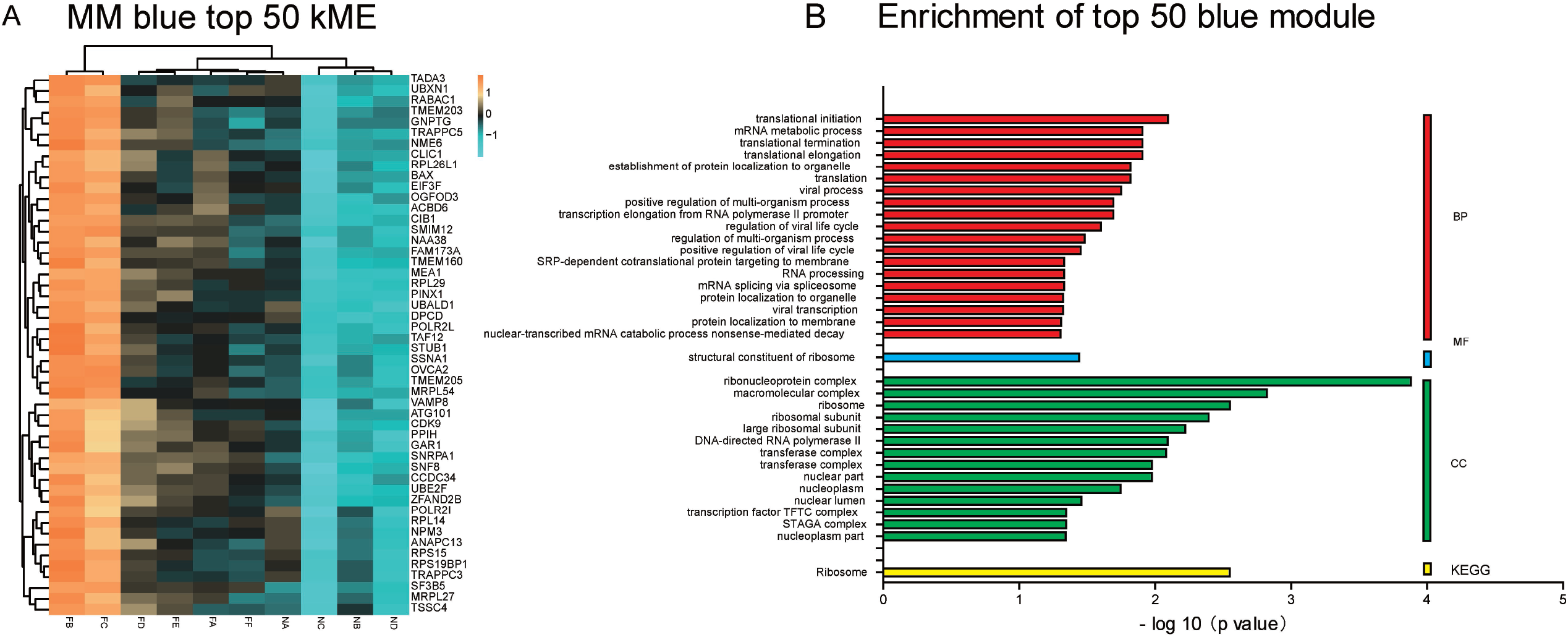
Blue module. A) Expression levels of the top 50 highly expressed genes in 3 groups. GO/KEGG analysis of the top 50 highly expressed genes.

**Figure 5.**
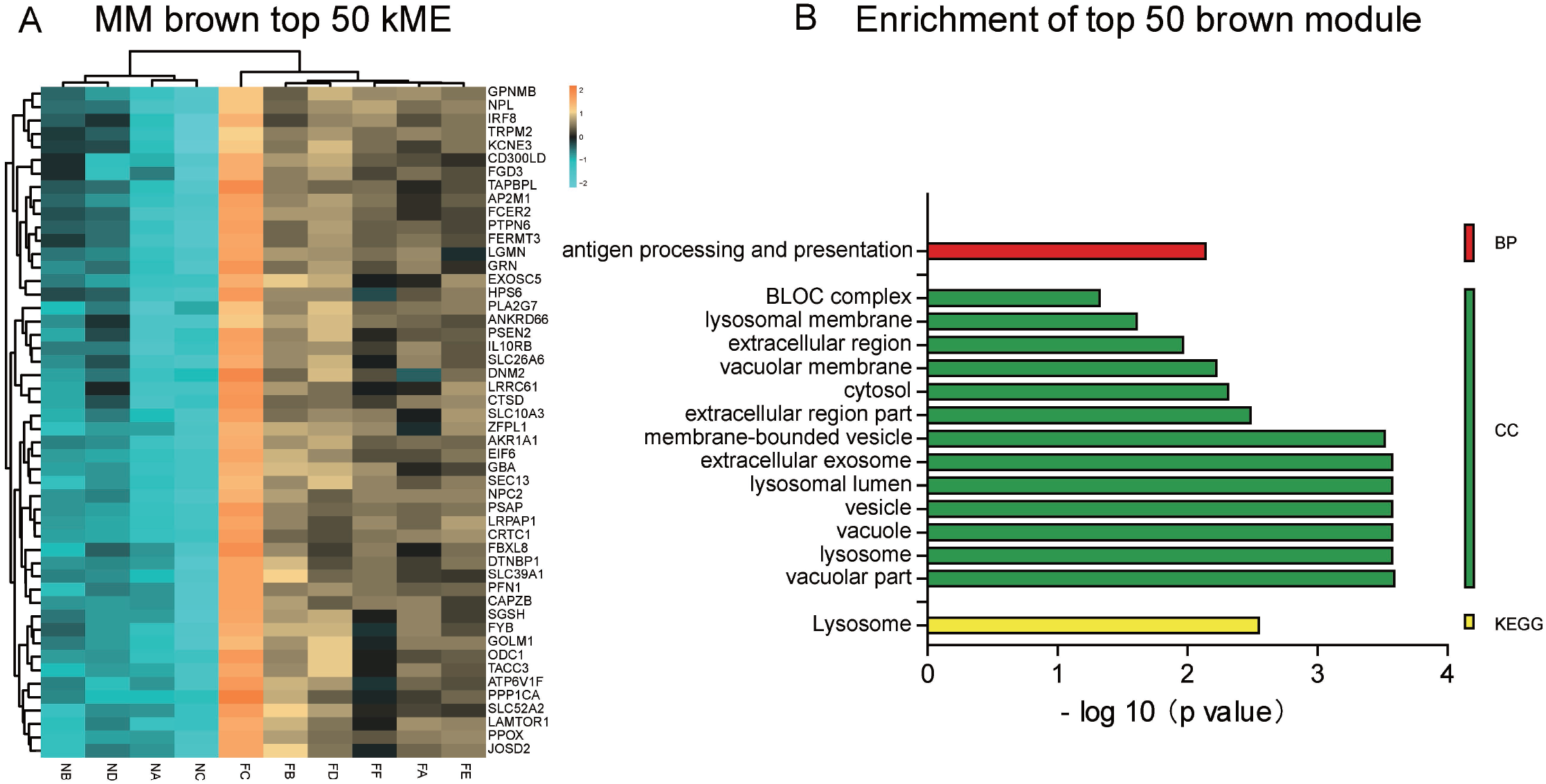
Brown module. A) Expression levels of the top 50 highly expressed genes in 3 groups. GO/KEGG analysis of the top 50 highly expressed genes.

### PPI regulatory network analysis of different modules

According to the data set from STRING, the PPI network consisted of 1775 gene signatures and 17,699 interactions based on 1861 DEGs in the three modules. A PPI regulatory network of the top 10 genes in degree in all three modules containing 30 nodes and 188 edges, including 20 up-regulated genes in the blue/brown modules and 10 down-regulated genes in the turquoise module, was constructed (Sup 1). The nodes with highest degrees were screened as the top 10 hub genes in each module. We further found that the top six genes with the highest degree in the blue module were AKT1, ATCB, SYK, PLEK, MMP9 and ITGAM, and the top six genes with the highest degree in the brown module were UBA52, NHP2L1, NHP2, UBC, RPL7A and RPS1. The interactions between each gene and its nearest neighbor are illustrated in Sup 2 and Sup 3.

AKT1, RPLP0, NEDD8, SSR4, SSR2, EXOSC4, LSM3, ARPC4 and CFL1 belonged to both blue and brown modules (intra-modular connectivity >0.9 in both modules) (Figure 6A). All nine shared genes are highly conserved in all species, including humans (Sup 4). We further identified AKT1 as the hub gene of this PPI network in the blue and brown modules using the same strategy according to node degree (Figure 6A). AKT1 and its relationship with eight other shared genes are illustrated in detail (Figure 6B).

**Figure 6.**
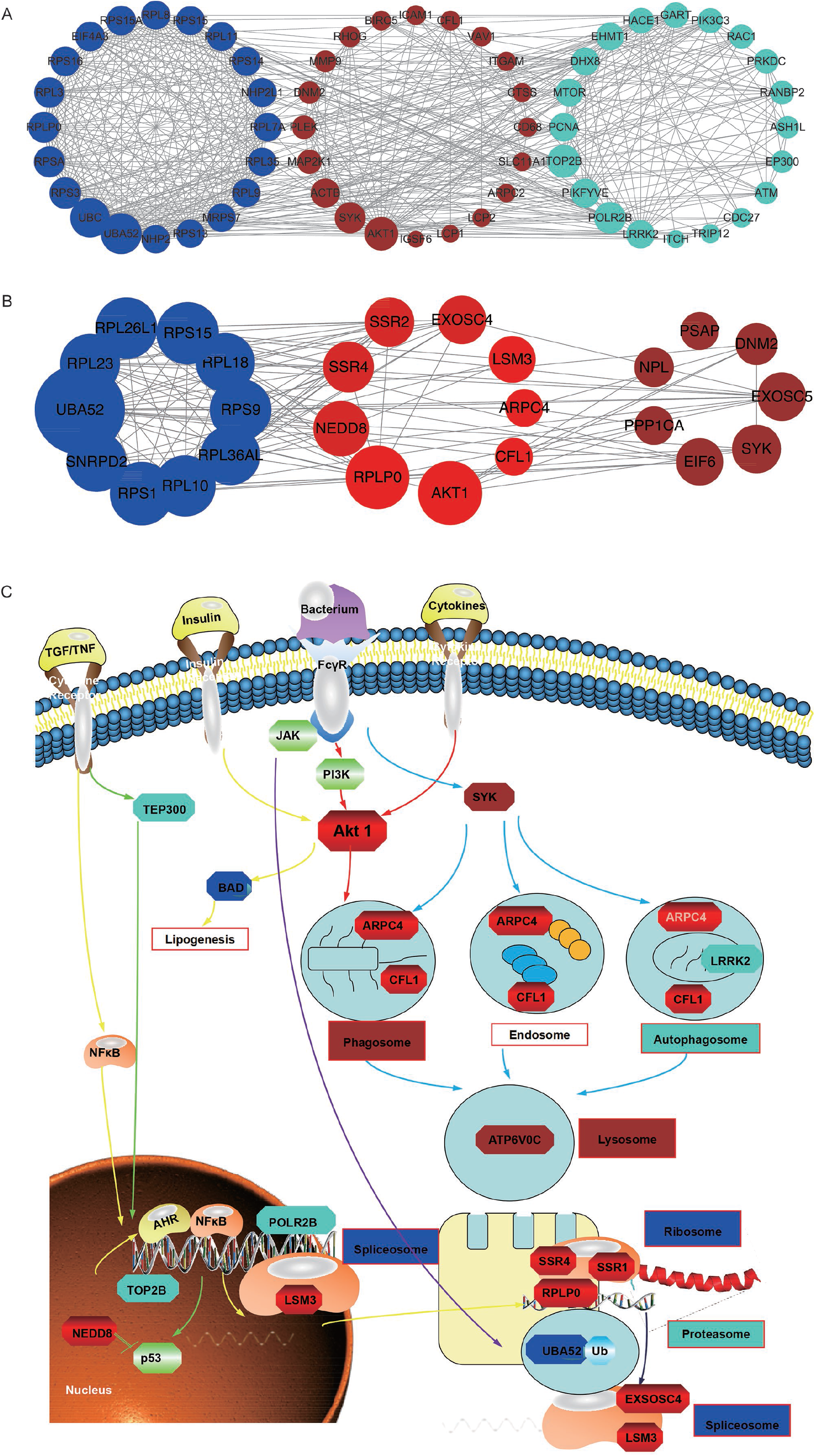
PPI regulatory network and gene relationships. A) Nine shared hub genes in both blue and brown modules. B) AKT1 and its relationship with eight other shared hub genes.

### The conservation of AKT1 between tree shrews and humans

Tree shrew AKT1 also consisted of an amino-terminal pleckstrin-homology domain, an inter-domain linker, a kinase domain and a 21 residue carboxy-terminal hydrophobic motif^36^. The comparison uncovered only 6 different amino acid residues in the functional structure, which were VAL-6, SER-31, SER-34, SER-148, ARG-170 and LYS-206. The three-dimensional domain structures of tree shrews and humans were dimensionally conserved (Figure 7A, 7B).

**Figure 7.**
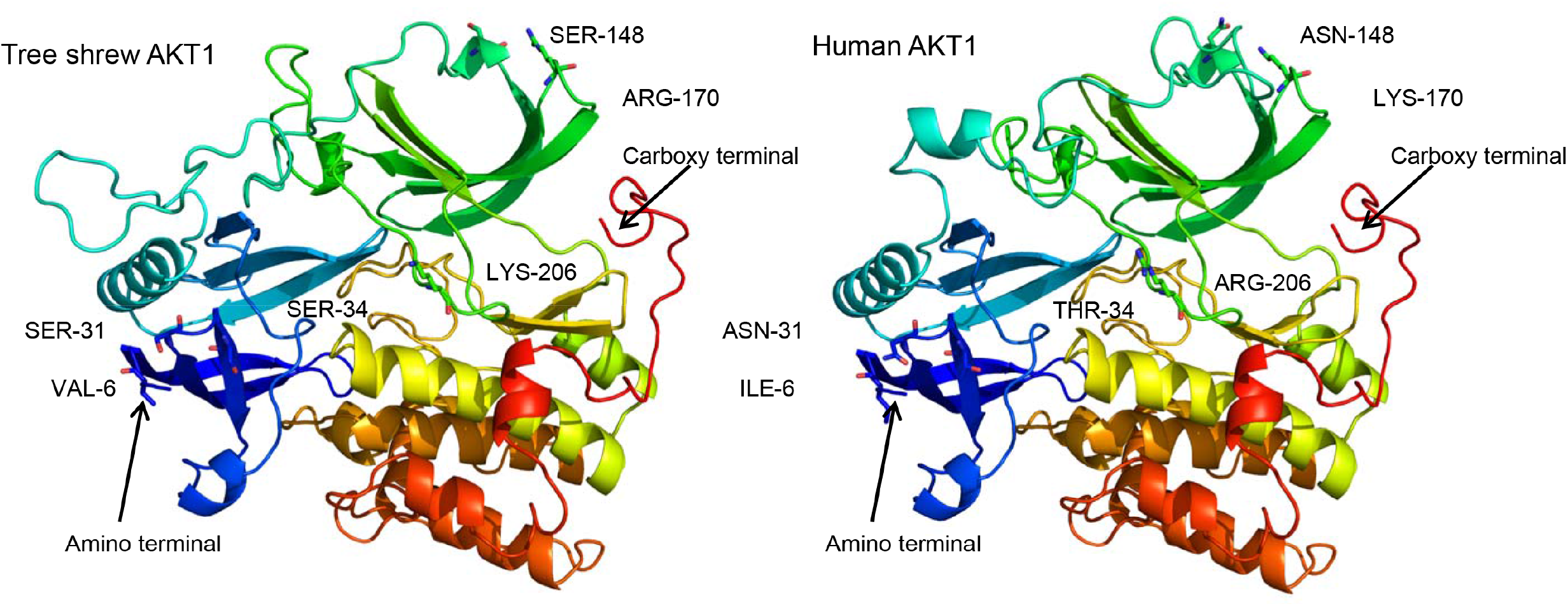
Conserved 3-dimensional structure of tree shrew and human AKT1.

### Validation of AKT1 and role of the PI3K/AKT pathway in obesity

We selected the hub gene AKT1 for certification. The AKT1 expression had a strong positive correlation with body weight (Figure 8A), Lee’s index (Figure 8B) and triglycerides (Figure 8D) and a relatively weak correlation with blood sugar (Figure 8C). The linear regression curves and statistics are shown (Figure 8).

**Figure 8.**
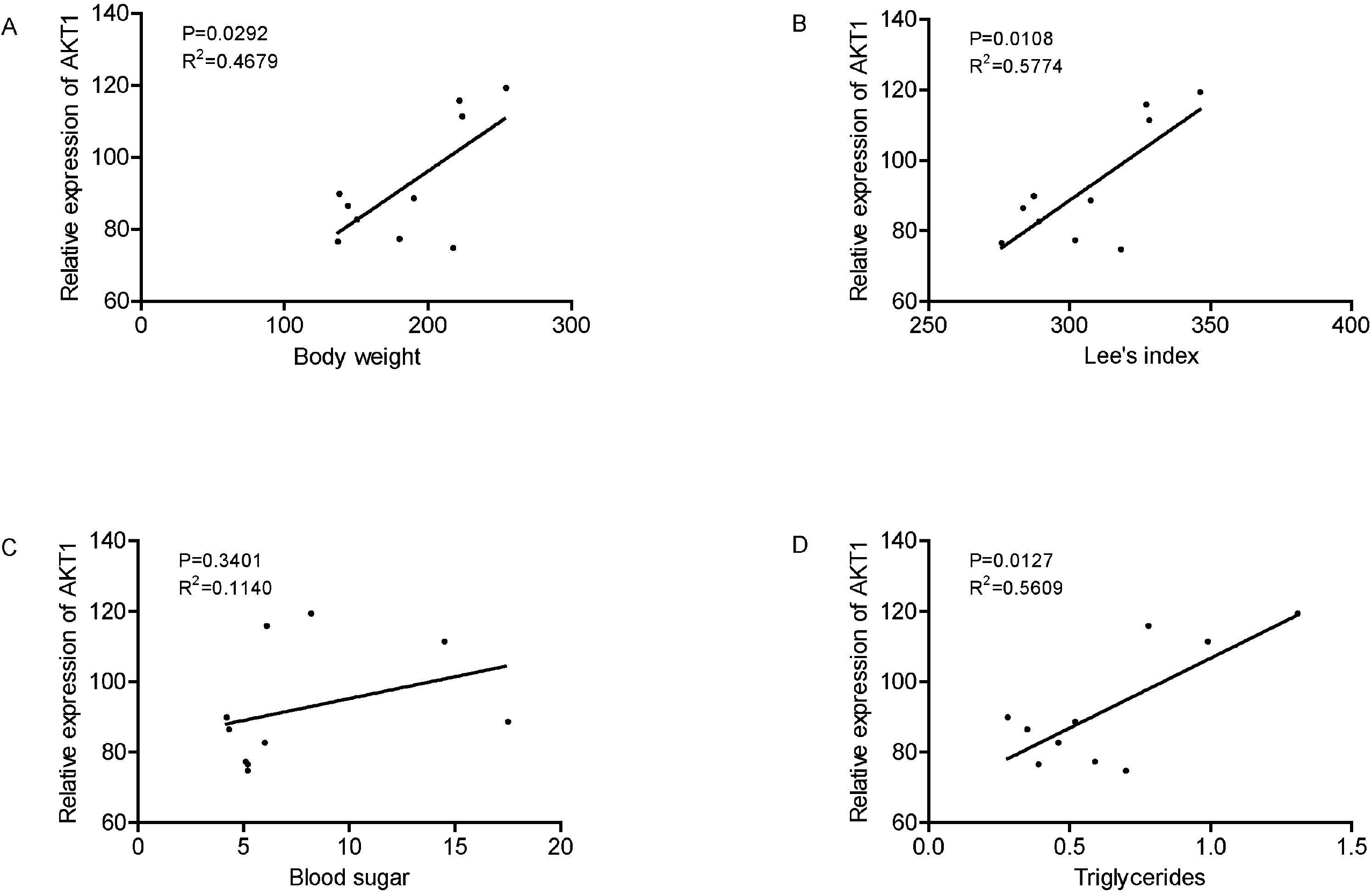
Validation of AKT1. (A) The correlation of AKT1 expression with body weight. (B) The correlation of AKT1 expression with Lee’s index. (C) The correlation of AKT1 expression with blood sugar. (D) The correlation of AKT1 expression with triglycerides.

When we focused on the AKT1 pathway, we found that 49 genes were differentially regulated (15 up-regulated and 34 down-regulated) in the PI3K-AKT pathway among the 3 modules. To further explore these genes, we performed a trend analysis with the Omicshare platform. All the changes described above are indicated by colors in the PI3K/AKT pathway: red for the 7 up-regulated genes, green for the 7 down-regulated genes, yellow for the 13 highly expressed genes and blue for the 6 genes with low expression in moderate obesity (Sup 5).

## Discussion

Obesity is a complicated metabolic and multifactorial status strongly associated with metabolic diseases, such as cancer, insulin resistance and cardiovascular disease ^37^, leading to public health burden and economic cost ^38^. Obesity in humans is classified according to the body mass index (BMI), which is divided into three categories: normal BMI (18.5–24.9 kg/m^2^), moderate obesity (25.0–34.9 kg/m^2^) and severe obesity (35.0–50.1 kg/m^2^)^39^. A BMI of 30 or higher in humans is currently classified as a disease^40^. To identify the co-expressed genes network for progression of obesity may provide an entry point for the molecular pathogenesis of obesity.

Notably, the obesity phenotypes were similar in our artificial breeding tree shrews. In this study, we characterized the spontaneous obesity phenotypes of tree shrew for the first time and explored the possible mechanism of obesity development based on the transcriptomic data. Like the phenotypes, the transcriptomic data of 10 tree shrews clustered in lean, moderate and severe fat groups. Clear between-group differences from RNA-Seq data were detected among the 3 groups.

WGCNA revealed some specific pathways that might play an important role in the progression of obesity (e.g., translational and metabolic pathways in the blue module, immune and stress-related pathways in the brown module and signal pathways in the turquoise module). Particularly, WGCNA revealed highly co-expressed genes in clusters.

Some of these genes were related to translation-relevant pathways in the blue module, e.g., the ‘ribosome’, ‘spliceosome’, ‘RNA polymerase’ and ‘protein processing in the ER’ pathways in the obese cohort. In obese animals, an increase in protein synthesis resulted in the increase in the ribosomal pathways and was accounting for the increase in energy demands^41^.

The brown module contained genes previously associated with immune function and stress, which confirmed the accepted close relation between obesity and other metabolic disorders, e.g., congenital disorders of metabolism (Padj = 4.80E-07), immune system diseases (Padj = 0.0001.3907-04) and cancers (Padj = 1.98E-03).

Because we focused mostly on up-regulated genes, we further identified AKT1, which belonged to both blue and brown modules, as the hub gene of this PPI network according to node degree. We also obtained the highest degree genes in each module, which were UBA52 in the blue module, AKT1 in the brown module and LRRK2 in the turquoise module.

UBA52 is the abbreviation of ubiquitin A-52 residue ribosomal protein fusion product 1. UBA52 not only supplies ubiquitin but also regulates the ribosomal protein complex. In previous studies, UBA52 was a key player in both ubiquitin-related cell cycle control and the ribosome complex in translational processes^42^. In the present study, obese tree shrews exhibited up-regulation of UBA52 at the mRNA level, suggesting its contribution to obesity possibly via interference with the cell cycle and the ribosome complex.

LRRK2 is the abbreviation for leucine-rich repeat kinase 2. Because it contains multiple conserved domains, the lrrk2 protein could interplay with many other proteins. In previous studies, LRRK2 kinase activity changes are associated with activation of the cellular death process and autophagy. Mutations in LRRK2 correlate with inherited and sporadic Parkinson’s disease. So, its contribution to obesity is possibly via interference with cell death and autophagy in accordance with cellular nutrient conditions^43^.

Akt1 is a conserved serine/threonine kinase that regulates and controls cell cycle progression, cell growth, cell metabolism and cell survival in many tissues and cell types. The 3-dimensional structure of AKT1 is highly conserved between tree shrews and humans (Figure 7). Chu et al. reported that overexpression of AKT1 promoted adipogenesis and led to lipoma formation in zebra fish^44^. Wan et al. reported that loss of AKT1 increased energy expenditure and protected against diet-induced obesity in rats^45^. In the present study, severe obese tree shrews exhibited up-regulation of AKT1 at the mRNA level, whereas moderate obese and lean tree shrews exhibited the same level of AKT1 mRNA expression, suggesting that AKT1 contributes to severe obesity development possibly via interference with signaling pathways and is a very important molecule linking severe obesity to cancer and diabetes^45^.

## Conclusion

System biology approaches are advantageous in unraveling transcriptional architecture of complex diseases^41^. We found the mRNA network, pathways and hub genes that were related to obesity. Furthermore, the tree shrew with different obesity phenotypes is a potentially useful model for studying the association of obesity with other diseases.

## List of tables

Sup table 1. Descriptive statistics (mean and standard deviation) and test of difference in means for a selection of obesity-related traits for the three subgroups.

Sup table 2. Overview of the most significantly overrepresented GO terms related to the modules discovered by WGCNA.

Sup table 3. Overview of the most significantly overrepresented KEGG pathways associated with the modules detected using WGCNA.

## List of figures

Figure 1

Figure 2

Figure 3

Figure 4

Figure 5

Figure 6

Figure 7

Figure 8

Sup 1

Sup 2

Sup 3

Sup 4

Sup 5

Sup 1 PPI regulatory network among the three modules.

Sup 2 Interactions among the top six hub genes and the nearest neighbors in the blue module.

Sup 3 Interactions among the top six hub genes and the nearest neighbors in the brown module.

Sup 4 Phylogenetic analysis of nine hub genes shared between the blue and brown modules shows a high degree of conservation.

Sup 5 AKT1 pathway–role of the PI3K/AKT pathway in obesity. Detailed expression pattern illustrated in the PI3K/AKT pathway.

## List of abbreviations

(WGCNA): weighted gene co-expression network analysis
(DEGs): differentially expressed genes
(WAT): white adipose tissue
(RNA-Seq): RNA-sequencing
(MMP): Maximal Mappable Prefix
(KEGG): Kyoto Encyclopedia of Genes and Genomes
(STRING): Search Tool for the Retrieval of Interacting Genes

## Declarations

### Ethics approval

This study was approved by theinstitutional Animal Care & Welfare Committee of IMBCAMS (NO. DWLL2016012).

### Consent for publication

Not applicable.

### Availability of data and material

The SRA (Sequence Read Archive) accession number for the exome sequences reported in this paper is SRP069216.

### Author’s contributions

YUANYUAN HAN completed the main experiment and wrote the main manuscript text; HUATANG ZHANG reviewed the manuscript; JIEJIE DAI and YANG CHEN provided the platform and ideas for this article.

## Acknowledgements

Not applicable.

